# Induced Disassembly of a Virus-Like Particle under Physiological Conditions for Venom Peptide Delivery

**DOI:** 10.1101/2020.09.01.278598

**Authors:** M. Patrick Kelly, Tanya Napolitano, Prachi Anand, Justin S. K. Ho, Shakeela Jabeen, Jessica Kuppan, Sujoy Manir, Mandë Holford

**Affiliations:** Department of Chemistry and Biochemistry, Hunter College, 695 Park Ave, New York, NY 10065, USA; The Ph.D. programs in Biochemistry, Chemistry and Biology Graduate Center of the City University of New York, 365 5^th^ Ave, New York, NY 10016, USA; Department of Invertebrate Zoology, The American Museum of Natural History, New York, NY 10024, USA; Department of Biochemistry, Weill Cornell Medicine, 413 E. 69^th^ Street, NY, NY 10021, USA

**Keywords:** Virus-like particle, P22 Bacteriophage, venom peptides, drug delivery, ROMP, induced disassembly

## Abstract

Virus-like nanoparticles (VLPs) show considerable promise for the *in vivo* delivery of therapeutic compounds such as bioactive venom peptides. While loading and targeting protocols have been developed for numerous VLP prototypes, induced disassembly under physiological conditions of neutral pH, moderate temperature, and aqueous medium, remain a challenge. Here, we implement and evaluate a ring-opening metathesis polymerization (ROMP) general mechanism for controllable VLP disassembly that is independent of cell-specific factors or the manipulation environmental conditions such as pH and temperature that cannot be readily controlled *in vivo*. The ROMP substrate norbornene is covalently conjugated to surface-exposed lysine residues of a P22 bacteriophage-derived VLP, and ROMP is induced by treatment of water-soluble ruthenium catalyst AquaMet. Disruption of the P22 shell and release of a GFP reporter is confirmed via native agarose electrophoresis and quantitative microscopy and light scattering analyses. Our ROMP disassembly strategy does not depend on the particular structure or morphology of the P22 nanocontainer and is adaptable to other VLP prototypes for the potential delivery of venom peptides for pharmacological applications.

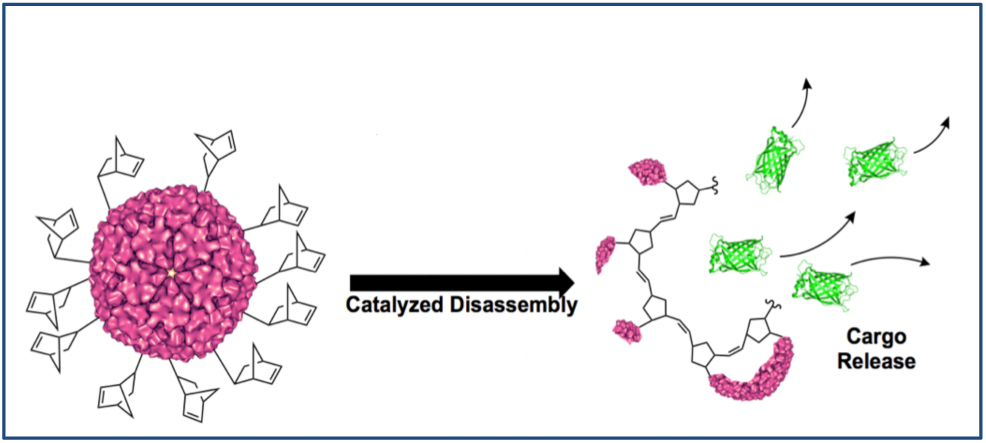

## Introduction

Venom peptides have immense therapeutic potential for the treatment of various human disease and disorders, including cancer and pain.^1–3^ As of 2018 there were more than 60 U.S. Food and Drug Administration-approved peptide drugs on the market, with hundreds more in various stages of development.^4–6^ Peptides as therapeutics are generally nontoxic, highly potent, and in most cases extremely specific, with few side effects.^7,8^ However, the delivery of therapeutic peptides to their molecular targets *in vivo* remains a significant challenge. Only a handful of peptides are robust enough to be administered orally, thus ruling out the most common and least invasive delivery route. Moreover, with the exception of specialized cell-penetrating peptides, very few peptides are able to cross the cell membrane or the blood-brain barrier (BBB).^9^ These hurdles have prevented the widespread development and application of venom peptides and bioactive peptides, in general. A prominent example of a venom peptide therapeutic that has enormous physiological potential but whose delivery hampers broad application is the drug ziconotide (Prialt®), which is derived from venomous cone snail *Conus magus* and used to treat chronic pain in HIV and cancer patients.^10,11^ Ziconotide does not cross the BBB and has to be administered by intrathecal injection. Despite the fact that ziconotide is a nonaddictive analgesic, active on N-type calcium channels and not opioid receptors, its invasive method of delivery restricts its use. There is a pressing need for innovative peptide drug delivery methods that will enable the untapped diversity of venom peptides to be utilized for pharmaceutical development.

One strategy for *in vivo* drug delivery is to package the therapeutic agent within a “Trojan Horse,” a macromolecular carrier that can protect the payload in transit and release it upon reaching the site of action. This approach has received significant attention recently for its role as a delivery mechanism for mRNA vaccines. For example, mRNA-1273, Moderna’s candidate vaccine for COVID-19, uses a lipid nanoparticle (LPN) to deliver an mRNA for the SARS-CoV-2 spike protein to cellular ribosomes.^12–14^ Other types of macromolecules have been studied for their potential as carriers as well, including natural and synthetic copolymers, inorganic particles, DNA origami, and noninfectious virus-like particles (VLPs) derived from the protein capsids of viruses.^15–20^ We previously used a VLP Trojan-Horse strategy to encapsulate ziconotide and shuttle it across a BBB model to demonstrate the proof-in-principle of venom peptide delivery using viral capids.^21,22^

Because viral capsids have specifically evolved to protect viral genomes and deliver them to host cells, VLPs are especially attractive candidates for macromolecular nanocontainers. VLPs can also be produced in large quantities through heterologous expression and precisely manipulated with the tools of protein engineering.^23^ Systems for the encapsulation of peptides or proteins have been reported for a number of VLP prototypes, including those derived from the cowpea chlorotic mottle virus (CCMV), Hepatitis B core antigen particles (HBVc), and the bacteriophages MS2, Qβ, and P22. These systems employ a variety of mechanisms for packaging peptides or proteins, including electrostatic interactions, passive diffusion, direct conjugation of the cargo to genomic RNA, and construction of a chimeric coat- or scaffold-cargo fusion.^24–28^

The bacteriophage P22 *Salmonella typhimurium* system used in this report allows for the sequestration of 200 – 300 copies of an arbitrary, genetically encoded protein cargo through the heterologous expression in *E. coli* of the P22 coat protein and a cargo-scaffold fusion protein.^28^ Co-expression of these two genes leads to self-assembly of P22 VLP nanocontainers loaded with the protein or peptide of interest, which can then be purified through standard methods. This system can produce very high effective local concentrations of the cargo package.^29^ In addition, to our method for translocating P22-derived VLPs containing the neuroactive ziconotide peptide across a BBB mimic, other strategies have also been reported for the cleavable release of cytotoxic cargo peptides in response to Cathepsin B, a protease overexpressed in many tumor cells.^21,22,30^ However, the development of a general, as opposed to cell-specific, mechanism for triggered VLP release *in vivo* remains an active area of research.

Here, we implement and evaluate a general mechanism for controllable disassembly that does not rely on either cell-specific factors or the manipulation environmental conditions such as pH and temperature that cannot be readily controlled *in vivo*.^31^ Our strategy for triggered disassembly of the P22 VLP nanocontainer employs the bioorthogonal ring-opening metathesis polymerization (ROMP) reaction, which is known to proceed under physiological conditions in the presence of a ruthenium catalyst (Grubbs Catalyst). ^32,33^ Our strategy consists of two steps: First, *N*-hydroxysuccinimide-activated norbornene moieties are covalently attached to multiple surface-exposed lysine residues on the VLP exterior via standard bioconjugation methods.^34^ Then, a bioorthogonal Ring-Opening Metathesis Polymerization (ROMP) reaction is initiated by the introduction of a water-tolerant ruthenium catalyst.^35,36^ The resulting ROMP reaction is driven by the release of ring strain in the norbornene substrate to trigger disassembly of the VLP and release of the cargo (Figure 1).

**Figure 1.**
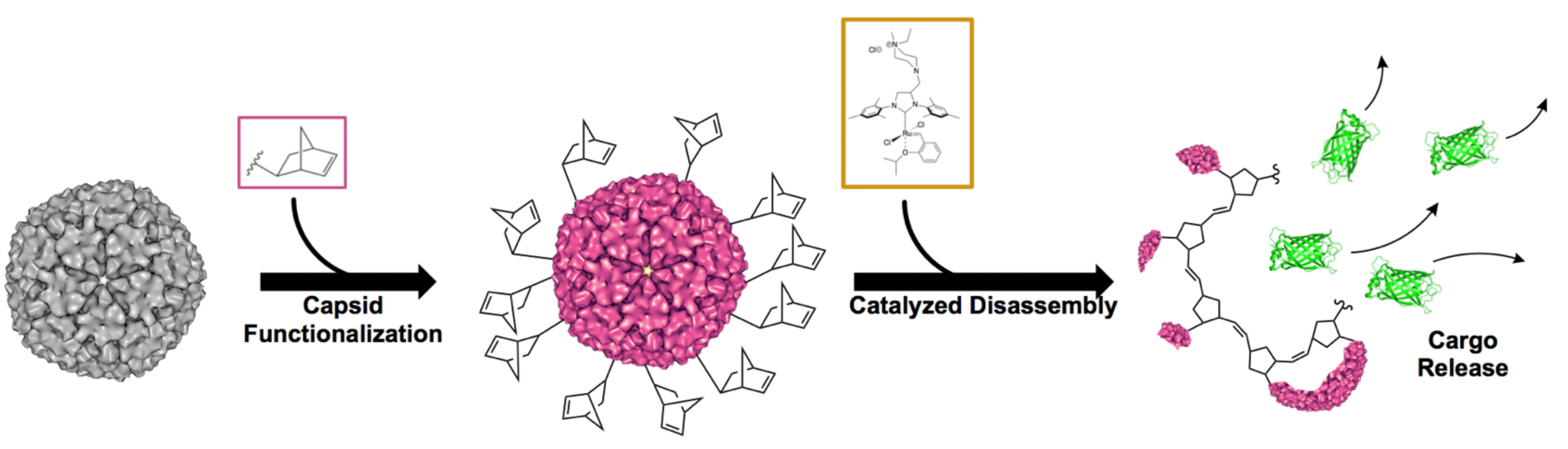
Physiological delivery of therapeutic peptides using functionalized VLP nanocontainers with a triggered disassembly strategy. First, P22-derived VLPs loaded with an arbitrary, genetically programmed cargo (here, GFP) are expressed and purified. Next, the loaded VLPs are functionalized by covalent attachment of a ROMP substrate (here, an NHS-activated norbornene-COOH) to surface exposed lysine residues. Finally, functionalized VLPs are disassembled under physiological conditions through the introduction of a ruthenium catalyst and initiation of Ring-Opening Metathesis Polymerization (ROMP).

ROMP, as the name suggests, is a polymerization reaction initiated by a transition metal catalyst and driven by the release of ring strain in a cyclic olefin such as cyclobutene, cyclopentene, ciscyclooctene or norbornene.^37^ The ROMP reaction is a powerful and broadly applicable tool for synthesizing macro-molecular substances.^38^ Its versatility in polymer chemistry has led to the development of well-defined ROMP catalysts with specialized properties, including catalysts with tailored initiation or propagation rates, water-tolerant and water-soluble catalysts, and photo-activated catalysts.^39–42^ The AquaMet catalyst utilized in this report is a variant of the widely-available Grubbs II ruthenium catalyst (Figure 2). The Grubbs catalysts are notable for their water-tolerance, for their effectiveness at low concentrations (as low as 2 mol%), and for the fact that the resulting ROMP reactions unfold on a physiologically relevant timescale (often on the order of minutes). In addition, the AquaMet catalyst incorporates a quaternary ammonium group and is thus highly water-soluble.^43,44^

**Figure 2.**
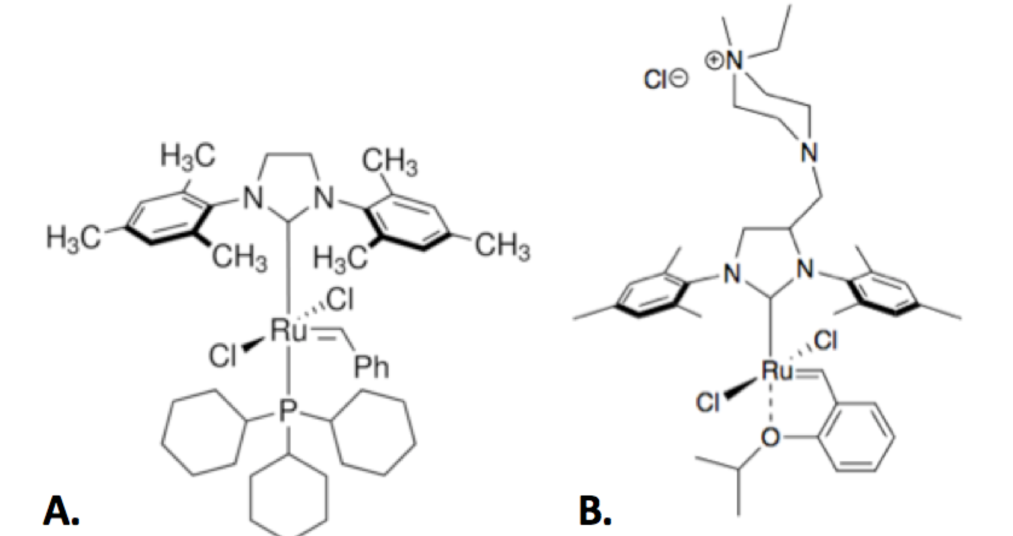
Ruthenium metathesis catalysts used for ROMP disassembly of VLP nanocontainers. The Grubbs II catalyst (A) and the AquaMet catalyst (B). The quaternary ammonium group appended to the N-heterocyclic carbine lig- and provides improved solubility in water.

In a previous study, we reported pilot results of this Trojan Horse strategy. P22-derived nanocontainers loaded with a GFP cargo (P22_GFP) were recombinantly expressed and purified, and the resulting VLP was functionalized by covalent conjugation of the ROMP substrate norbornene to the capsid surface. Treatment of the functionalized nanocontainers with a second-generation Grubbs catalyst (Grubbs II) resulted in significant morphological distortion with respect to untreated constructs, as indicated by TEM micrographs. However, direct evidence of cargo release was not observed.^22^ In the present study, we reexamine the induced disassembly of norbornene-functionalized P22_GFP constructs treated with the water-soluble AquaMet catalyst (Figure 2).^45^ In addition, a cargo-release assay was developed and implemented to monitor the release of the packaged GFP reporter into the solvent phase following catalyst treatment.

## Results and Discussion

To determine the catalyst concentration needed to trigger the ROMP reaction and subsequent disassembly in functionalized capsids, serial dilutions of catalyst were prepared and used to treat aliquots of P22 nanocontainers. To monitor for non-specific catalyst effects, nonfunctionalized P22 nanocontainers were used as controls. Reactions were incubated overnight at room temperature and characterized by native agarose electrophoresis. The resulting native agarose gels indicate a clear difference in the behavior of nanocontainers conjugated with the ROMP-substrate norbornene (P22His_6_GFP_Norb) and nano-containers that lack the norbornene moiety (P22His_6_GFP) (Figure 3A). A non-specific catalyst effect is observed at high catalyst concentrations (Lane 5), however, functionalized nanocontainers (P22His_6_GFP_Norb) appear to dissociate at catalyst concentrations that have no observable effect on non-conjugated nanocontainers (P22His_6_GFP_Norb).

**Figure 3.**
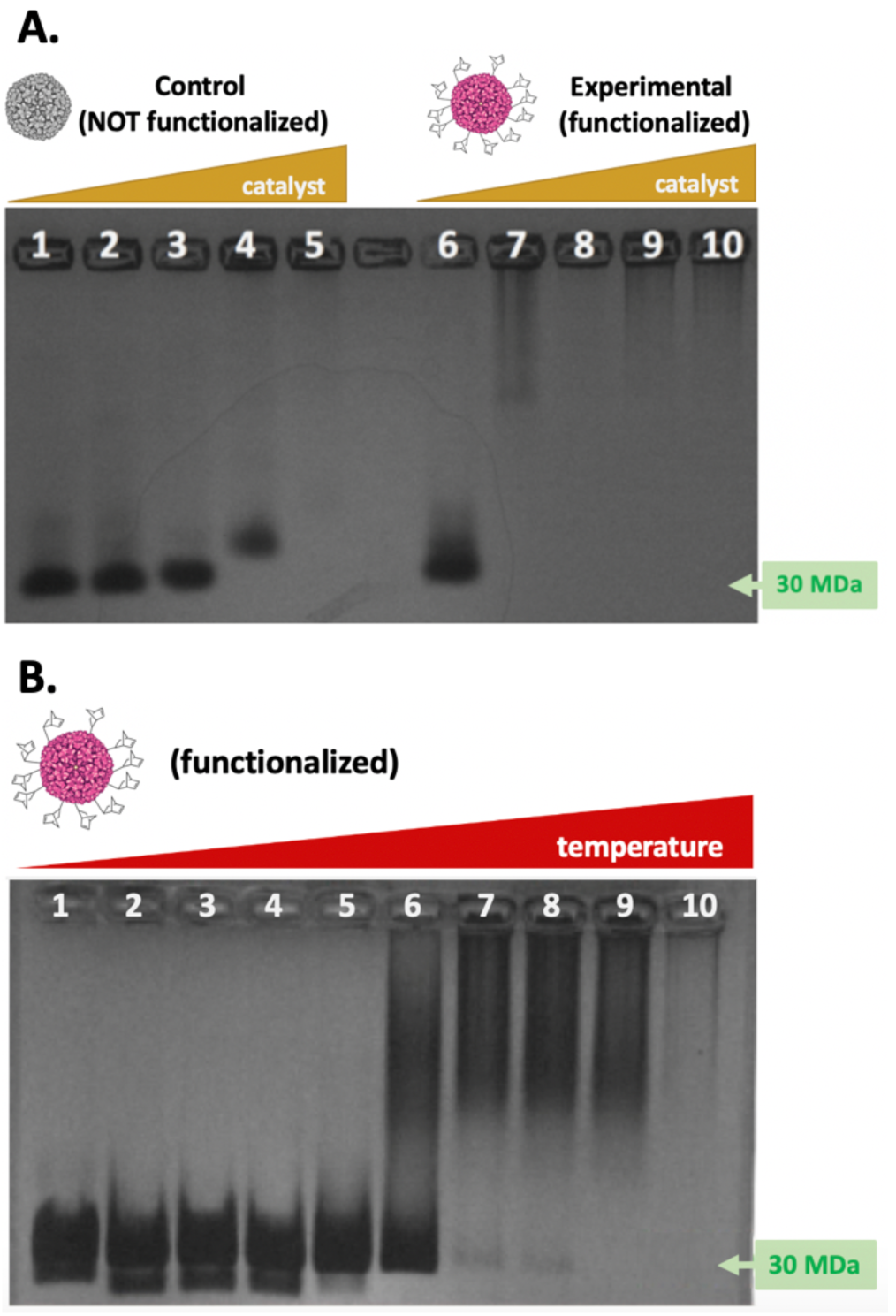
Catalyst-activated disassembly of P22-derived VLP nanocontainers. (A) Treatment of 70 μL (1.29 mg/mL) aliquots of nonfunctionalized (P22His6GFP_WT, left) and functionalized (P22His6GFP_Norb, right) nanocontainers with increasing concentrations (0 - 0.28 mg/mL) of AquaMet catalyst indicates capsid disassembly at significantly lower catalyst concentrations for functionalized nanocontainers. While functionalized capsids disassemble at 0.054 mg/mL catalyst (Lane 7), the 30 MDa band corresponding to the fully assembled capsid only disappears at a catalyst concentration of 0.28 mg/mL (Lane 5) for nonfunctionalized capsids. (B) Heat-activated disassembly of functionalized nanocontainers (P22GFP_Norb). Functionalized nanocontainers (P22_GFP_Norb) were divided into 15 μL aliquots and heated for 10m at a temperature gradient of 50°C - 85°C, then subject to native agarose electrophoresis. The on- set of disassembly is observed in the 65°C - 70°C range (lanes 6 - 7), as indicated by smearing and gradual disappearance of the 30MDa band visible in the control (Lane 1, no heat). Note that band smearing is also visible in Lanes 7 - 10 in (A) above.

We next compared ROMP-induced disassembly with heat-induced disassembly. Past studies have demonstrated that heating P22 VLPs at 65°C results in capsid expansion, a morphological change that mimics the expansion of capsids during DNA packaging in the natural P22 life cycle. Additional heating at 75°C induces release of the pentameric subunits from the fivefold icosahedral vertices, resulting in the so-called “wiffle-ball” conformation and the release of packaged cargo.^46^ To examine the behavior of functionalized capsids under a thermal gradient, we again used native agarose electrophoresis (Figure 3B). As expected, heat-induced disassembly was observed, albeit at a slightly lower temperature (starting at 70°C) than reported for the case of non-functionalized capsids (>80°C). Importantly, treatment of functionalized P22 constructs with catalyst concentrations above 0.054 mg/mL produced smeared bands (Figure 3A, Lanes 7-10) that resemble those of capsids heated at 65°C or higher (Figure 3B, Lanes 6 – 10).

Dynamic Light Scattering (DLS) confirmed that ROMP functionalization did not significantly alter either the size or the morphology, as measured by the hydrodynamic diameter and polydispersity index (PDI), of P22 nanocontainers. However, DLS analysis of functionalized nanocontainers (P22His_6_GFP_Norb) treated with AquaMet catalyst revealed a pronounced, dose-dependent increase in both the mean hydrodynamic diameter and the PDI with increasing catalyst concentration (Figure 4). These results are consistent with ROMP-induced dissociation and/or aggregation.

**Figure 4.**
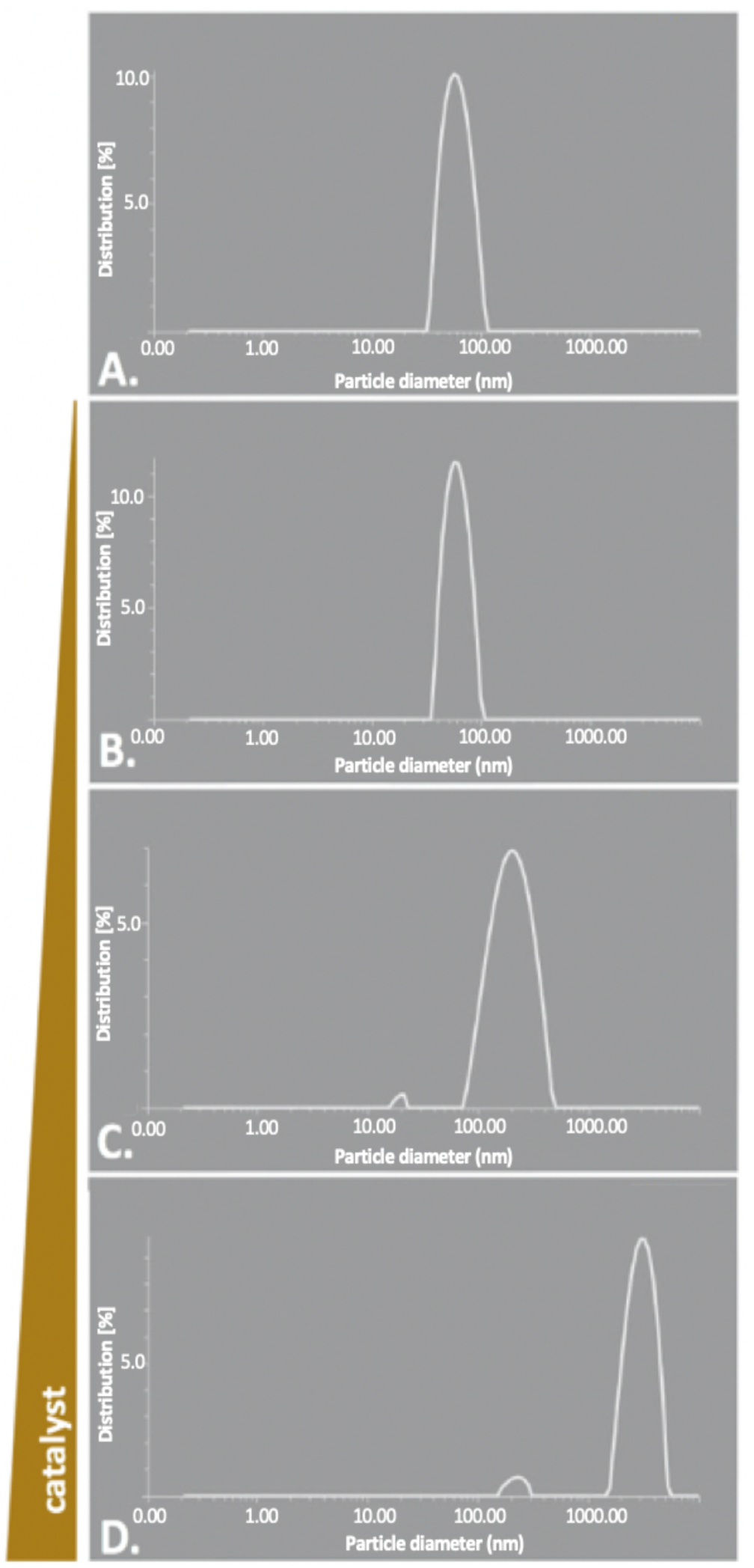
DLS characterization of norbornene-conjugated nanocontainers. Functionalized nanocontainers (P22His_6_GFP_Norb) were divided into 82.5 μL aliquots (1.09 mg/mL) and treated with increasing concentrations (0 - 0.325 mg/mL) of catalyst. DLS characterization of the hydrodynamic diameter and polydispersity index (PDI) reveals a steady migration of the size distribution peak towards larger diameters accompanied by an increase in PDI with increasing catalyst concentration. (A): no catalyst, dmean = 60.2 nm, PDI = 15.1%. (B): 13 μg/mL catalyst, dmean = 60.6 nm, PDI = 7.5%. (C): 130 μg/mL catalyst, dmean = 193.5 nm, PDI = 21.9%. (D): 325 μg/mL catalyst, dmean = 3222 nm, PDI = 26.9%.

TEM micrographs of norbornene-conjugated nanocontainers (P22His_6_GFP_Norb) treated with AquaMet water-soluble ruthenium catalyst exhibit significant distortion in capsid morphology. Aggregates of what appear to be disrupted and fused capsids are plainly visible. We were unable to locate any of the regular icosahedral structures characteristic of P22 VLP not reacted with the AquaMet catalyst (Figure 5). The observation of disordered aggregates is consistent with a polymerization reaction that occurs at both intra- and inter-nanocontainer inter-faces.

**Figure 5:**
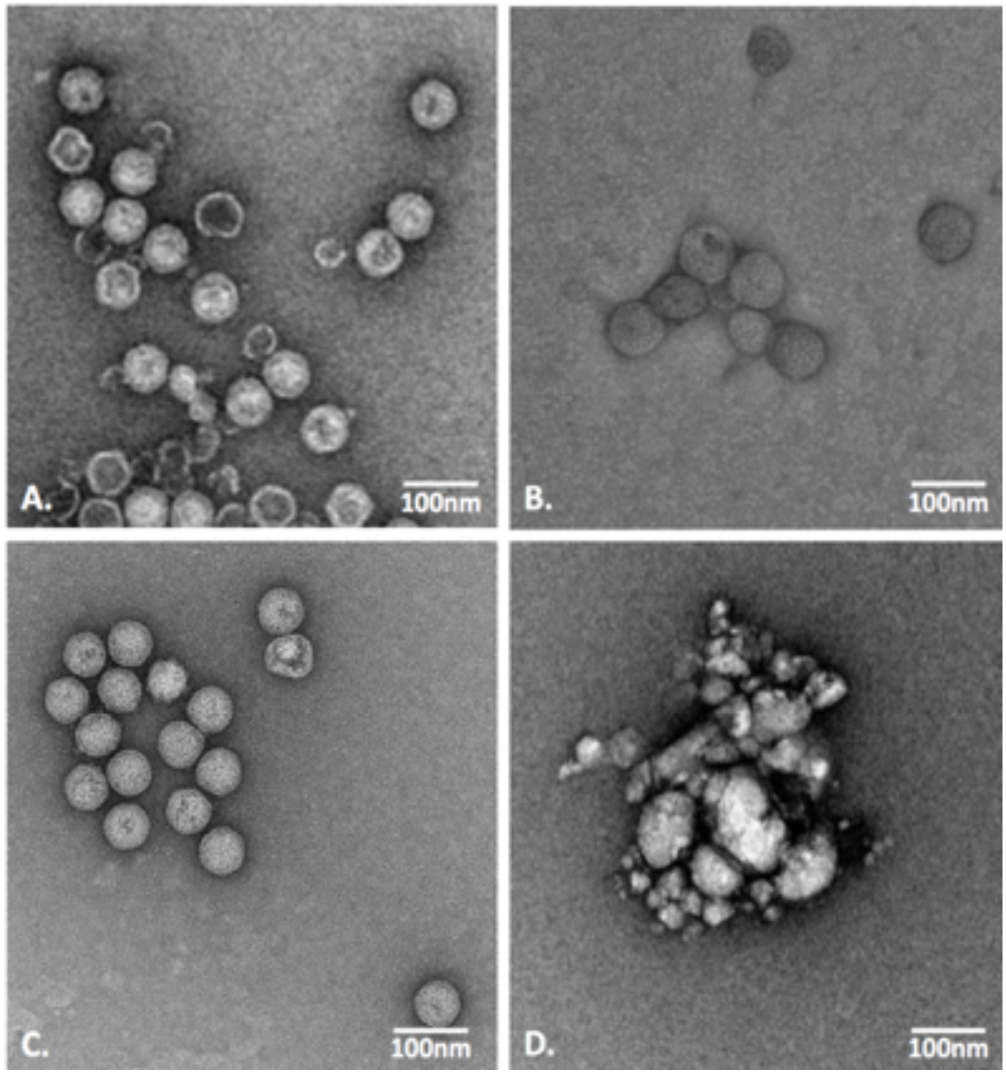
Representative TEM images. Images show (A) untreated, nonfunctionalized constructs (P22His_6_GFP_WT, no catalyst), (B) untreated, functionalized constructs (P22His_6_GFP_Norb, no catalyst), (C) nonfunctionalized constructs treated with AquaMet catalyst (P22His_6_GFP_WT, 0.45 mg/ml catalyst), and (D) functionalized constructs treated with AquaMet catalyst (P22His_6_GFP_Norb, 0.325 mg/ml catalyst).

To further confirm that P22 ROMP functionalized nanocontainers were undergoing structural disruptions that may indicate disassembly, we developed a cargo release fluorescence assay in which we monitored the release of the His_6_-GFP cargo from control (nonfunctionalized, P22His_6_GFP) and experimental (functionalized, P22His_6_GFP_Norb) constructs treated with 0.212 mg catalyst. After treatment, each of the samples was incubated with Ni^2+^ Sepharose beads then washed and eluted, as described below. The four wash fractions and four elution fractions were monitored for absorbance at 280 nm (aromatic amino acids λ_max_) and 495 nm (GFP λ_max_). A significant peak was observed in the elution fractions of the experimental sample at both wavelengths (Figure 6). No peak was observed in the elution fractions of the control sample. These results demonstrate induced release of the cargo protein at near-physiological conditions (PBS, 25°C).

**Figure 6.**
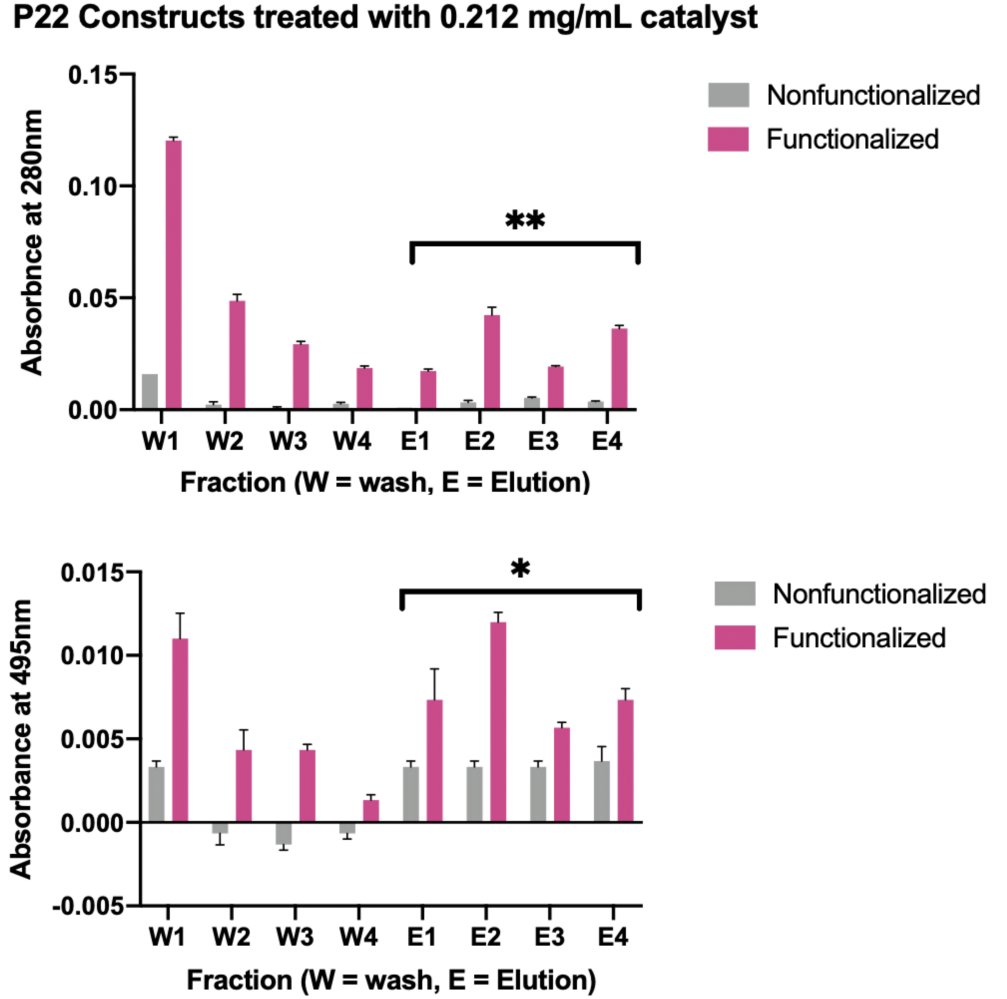
Cargo release assay indicate induced release of GFP protein at near-physiological conditions. Functionalized (magenta) and nonfunctionalized (grey) constructs were loaded with 200-300 copies of a His-tagged GFP reporter and treated with 0.212 mg/mL AquaMet catalyst. Samples were incubated with Ni^2+^ Sepharose beads, then washed and eluted. Absorbance profiles of functionalized constructs exhibit a strong peak at the start of elution (fraction E1) for both 280 nm and 495 nm (the λ_max_ of GFP) with p** = 0.066 at 280 nm and p* = 0.0142 at 495 nm for functionalized vs. nonfunctionalized elution fractions, respectively.

To monitor cytotoxicity of the nonfunctionalized P22His_6_GFP and functionalized P22His_6_GFP_Norb constructs, MTT assays were conducted using BJ normal foreskin epithelial cells. No evidence of cytotoxicity was found for treatment with VLP constructs at concentrations up to 100 μg/mL (Figure 7A).

**Figure 7.**
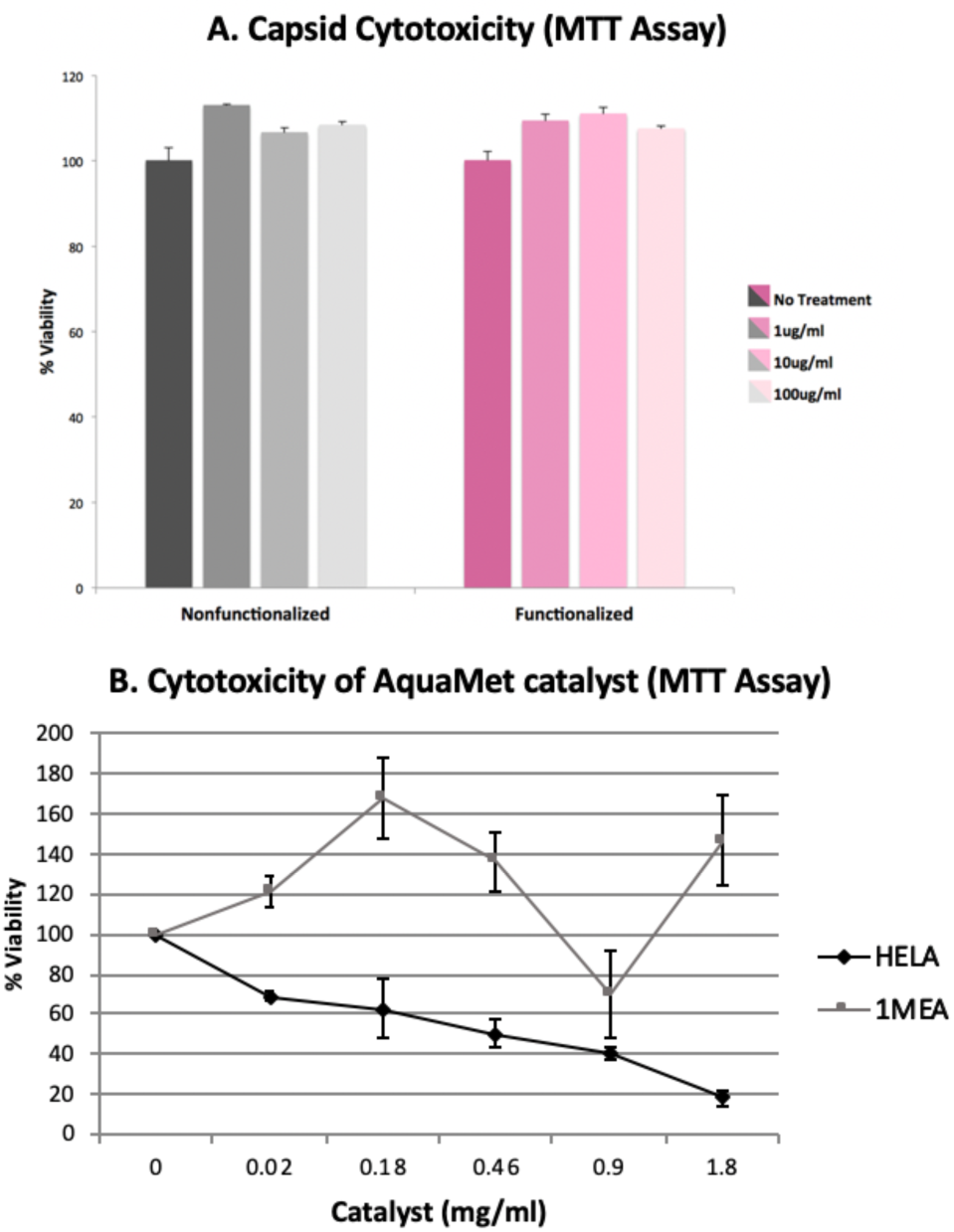
Cytotoxicity of P22 constructs and ruthenium catalyst. (A) Nonfunctionalized P22His_6_GFP and functionalized P22His_6_GFP_Norb capsids are not cytotoxic. In MTT assays using BJ normal foreskin epithelial cells, neither nonfunctionalized (grey) nor functionalized (magenta) P22 VLP constructs show significant cytotoxicity below 0.1 mg/mL. (B) In MTT assays using 1MEA and HeLa cell lines with varying concentrations of AquaMet catalyst, it was found that the catalyst is moderately cytotoxic in HeLa cells at concentrations above ∼0.2 mg/mL which is well above what is required for ROMP reaction to induce disruption in the VLP.

Finally, to assess the cytotoxicity of the AquaMet catalyst, we conducted MTT assays in HeLa and 1MEA cell lines. While moderate cytotoxicity (>10% cytotoxicity, <60% viability) was found for catalyst concentrations above 200 μg/mL in HeLa cells, this value is significantly higher than the 50-100 μg/mL needed to disrupt functionalized P22His_6_GFP_Norb constructs without affecting nonfunctionalized P22His_6_GFP constructs (Figure 7B). As a reference, these catalyst concentrations are roughly comparable to the reported peak serum concentrations of a variety of common drugs, including ibuprofen (61.1 ± 5.5 μg/mL), aspirin (24 ± 4 μg/mL), the antiviral cidofovir (19.6 ± 7.2 μg/mL), and the β-lactam antibiotic cefepime (65 ± 7 μg/mL).^47^ Finally, no significant cytotoxicity was found for 1MEA cells.

## Conclusions

Native agarose electrophoresis, dynamic light scattering (DLS), and transmission electron microscopy (TEM) all indicate that treatment of ROMP-functionalized P22 nanocontainers with a water-soluble ruthenium catalyst (AquaMet) results in significant disassembly of the P22 constructs. Studies of the physical virology of the P22 bacteriophage confirm that the ROMP reaction induced here provides sufficient energy for disassembly. For example, one study estimated that each of the 420 coat protein monomer contributes -7.2 kcal/mol to procapsid stability, while each of the 200-300 scaffold proteins contributes -6.1 kcal/mol.^48^ These estimates assume that 1) each scaffold protein contributes the whole of its binding energy to pro-capsid stability, and 2) that all procapsids contain the same number of scaffold proteins.^49^ While these assumptions are not entirely realistic, they provide an upper-bound estimate for the binding energy of the procapsid as a whole: (420 coat proteins)(−7.2 kcal/mol) + (300 scaffold proteins)(−6.1 kcal/mol) ≈ 5000 kcal/mol. As noted above, polymer elongation by ROMP is driven by the release of ring strain in a cyclic olefin substrate. The ring strain of norbornene, the ROMP substrate used here, is 27.2 kcal/mol.^50^ With an average of 4.12 norbornene-functionalized lysines per coat protein monomer, this yields approximately 47,000 kcal/mol ring strain energy per functionalized capsid.^51^ Thus, only ∼10% of the theoretical yield of norbornene ring strain energy needs to be harnessed to reach the disassembly threshold.

We also found significant evidence of cargo release under physiological conditions, as indicated by an increase in absorbance at 280 nm and 495 nm following capsid disruption. The increase at the 495 nm peak in particular indicates that the released cargo was still functional, and thus that the tertiary structure was not significantly altered. Finally, the catalyst concentrations needed for capsid disruption are not significantly cytotoxic and are within the effective concentrations of some common pharmaceuticals. VLPs and VLP-derived nanoparticles have been studied extensively in recent years for their biomedical potential, in particular for use in gene therapy and the targeted delivery of therapeutic and diagnostic agents.^20,52^ While gene therapy applications generally use VLPs derived from mammalian viruses, therapeutic and diagnostic applications tend to focus on plant viruses and bacteriophages, which are less likely to trigger pathogenic virus-host interactions.^53^ One recent study found no evidence of overt toxicity in naïve and immunized mice after single injections of protein cages derived from the Cowpea Chlorotic Mottle Virus (CCMV) and the heat shock protein cage (Hsp). Both CCMV and Hsp exhibited broad distribution throughout most tissues and organs, and were rapidly excreted, with no evidence of long-term persistence.^54^ While these findings suggest that P22-derived nanocontainers may be safe for biomedical applications, a comprehensive study has yet to be conducted.^54^ Additionally, while the potential toxicity of the transition-metal olefin catalysts has not been well researched, there are some indications that ruthenium (Ru) is genotoxic.^55^ However, the polynorbornene macromolecule that results from norbornene polymerization is not cytotoxic.^56^ These concerns are offset by the fact that very small quantities of ruthenium generally suffice to catalyze ROMP.^57^

In contrast to other VLP delivery systems, which rely on passive diffusion or environmental conditions such as hydrophobicity,^58^ pH (typically through the endosomal pathway), and cell-specific proteases^30^ for cargo release, the system described here employs a bioorthogonal ROMP reaction. Importantly, the ROMP delivery system is modular: First, the protein cargo is genetically encoded, and thus arbitrary. Further, the functionalization method (bioconjugation via an activated NHS-ester) and trigger protocol (treatment with a water-soluble ruthenium catalyst) are completely general, and expected to translate to other VLP systems, whether derived from other phage particles or wholly synthetic. Finally, the ROMP reaction itself is bioorthogonal, and is thus independent of environmental conditions and cell-specific factors.

While surface-exposed lysine residues were used as functionalization handles in this report, selective bioconjugation is also possible on cysteine, tyrosine, and methionine residues.^59^ Future efforts might thus focus on the construction of bifunctional VLPs decorated with targeting molecules such as cell-penetrating peptides or antibody fragments in addition to ROMP substrates. We envision a wholly modular peptide drug delivery system that combines a genetically-programmed cargo with an arbitrary targeting moiety and an interchangeable VLP chassis. Such a drug delivery vehicle would significantly advance the use of venom peptides for therapeutic development.

## Experimental Procedures

### Construction of P22His_6_GFP nanocontainers

A Q5 sitedirected mutagenesis kit (New England Biolabs) was used to insert a hexahistidine tag at the N-terminal of the SP141 gene of the P22GFP expression plasmid via standard PCR protocols. Transformed clones were screened for correct insertion of the hexahistidine tag and confirmed by Sanger sequencing (Genewiz, Figure S1). P22His_6_GFP Nanocontainers were expressed and purified as described in O’Neil et al. Briefly, BL21(DE3) E. coli were transformed with the P22His_6_GFP expression plasmid, grown to an OD600 = 0.6 in LB medium and induced with isopropyl β-D-1-thiogalactopyranoside (IPTG, 1mM). After 4 hours, cells were harvested, lysed by treatment with DNase, RNase, and lysozyme, then sonicated. The clarified lysate was ultracentrifuged over a 35% (w/v) sucrose cushion, dialyzed against pH 7.2 PBS overnight, then concentrated and resuspended in pH 7.2 PBS using a 100kDa MWCO centrifugal filter device (Amplicon). Expression and self-assembly of P22 VLPs was confimed by native agarose electrophoresis (Figure 3A) and ESI-MS (Figure S5).

### Conjugation of norbornene to P22His_6_GFP VLPs

To conjugate norbornene to surface-exposed lysine residues on the P22 bacteriophage capsid, 19.2 mg (1 x 10-4 mol) of 1-ethyl-3-(3-dimethylaminopropyl) carbodiimide (EDC) and 21.2 mg (1 x 10-4 mol) N-hydroxysuccinimide (sulfo-NHS) were added to 1 mL of a 0.1M solution of 5-norbornene-2-carboxylic acid in 0.1M phosphate buffer (pH 6.0) (all reagents purchased from Sigma-Aldrich). The solution was allowed to react at room temperature with intermittent vortexing. After 30m, 180 μL of the sulfo-NHS/EDC/Norbornene solution was added to 1 mL of P22His_6_GFP (5 mg/mL) in PBS (pH 7.6). After reacting at room temperature overnight on a shaker at 250 rpm, the product was concentrated by means of a 100 KDa MWCO centrifugal filter device (Amplicon) and resuspended in pH 7.2 phosphate buffer. Conjugation of norbornene-COOH to the P22 coat protein was confirmed by ESI-MS (Figure S6).

### DLS characterization of P22His_6_GFP nanocontainers

P22His_6_GFP nanocontainers at a concentration of 10μM in Ph 7.2 PBS were filtered through a 0.2 μm syringe filter and transferred to a low-volume (45 μL) quartz cuvette for DLS analysis with an Antor Paar Litesizer 500 instrument. All data were analyzed using the Kalliope software package.

### Native agarose electrophoresis

All P22His_6_GFP_Norb samples were normalized with respect to protein concentration prior to native agarose electrophoresis. A 20μL aliquot was removed from each sample, mixed with loading buffer (40% glycerol, bromophenol blue) and loaded into the wells of a 1.0% native agarose gel. Gels were run at 65V for 2.5 hours in TBE buffer (89mM Tris, 89mM borate, 2mM EDTA), then stained with coomassie blue and distained with acetic acid and methanol. Images were produced using a Foto/Analyst FX imaging system (Fotodyne).

### Cargo-release disassembly assay

250 μL aliquots of a 2.5mg/mL solution of P22His_6_GFP_Norb in PBS were placed in each of two microcentrifuge tubes. To the experimental sample was added 66.4 μL of a 3.2 mg/mL solution of AquaMet catalyst in ddH_2_O (final concentration 212 μg/mL). To the control sample was added 66.4 μL of ddH_2_O. Samples were incubated at room temperature overnight, then supplemented with 250 μL wash buffer (Tris-HCl (10 mM), imidazole (20 mM), NaCl (200 mM), pH 8) and 50 μL Ni^2+^ Sepharose beads and placed on a nutator. After 1h, the beads were collected by centrifugation (1000g, 1m) and the flow through removed. Beads were resuspended four times in 100 μL of wash buffer, then eluted four times with 100 μL wash buffer supplemented with 200 mM imidazole. Wash and elution fractions were monitored for absorbance at 280 nm and 495 nm (peak absorbance of GFP) on a NanoDrop 2000c spectrophotometer (Thermo Scientific).

### MTT cytotoxicity assay: VLP constructs

The cytotoxicity of VLPs was examined in BJ normal human foreskin epithelial fibroblasts by MTT assay. 5000 cells were seeded in each well of a 96-well plate and incubated with P22His_6_GFP and P22His_6_GFP_Norb in quadruplicate at three concentrations: 1 μg/ml, 10 μg/ml, and 100 μg/ml. After 24h incubation, 20 μl of MTT solution (5.0 mg/ml methyl tetrazolium salt in PBS) was added to each well and plates were incubated for an additional 3h at 37°C in 5% CO2. After 3h, medium supplemented with MTT solution was aspirated and 100 μl of acidified isopropanol was added to each well. Spectrophotometric assays were conducted in a PowerWave HT Microplate Spectrophotometer at 550 nm and 620 nm. The mean and standard deviation of the δOD values were calculated by subtracting the 620 nm values from the 550 nm values. The absorbance of control cells (untreated) was used as the 100% viability baseline.

### MTT Cytotoxicity Assay: ROMP Catalyst

The cytotoxicity of AquaMet catalyst was examined in HeLa and 1MEA cells by MTT assay, as described above. The cell lines BNL1MEA.7R.1, also called 1MEA (mouse liver carcinoma), was a kind gift from Professor Olorenseun Ogunwobi (Hunter College, New York, NY). Cells were maintained in Dulbecco’s modified Eagle’s medium (DMEM, Gibco-BRL Life Technologies, Paisley, UK) supplemented with 10% fetal bovine serum (FBS), 1% penicillin, 4.5g/L glucose, L-glutamine, and sodium pyruvate. HeLa (human cervical cancer, American Type Culture Collection, Rockville, MD), were regularly cultured in Eagle’s minimal essential medium (Gibco Co., Grand Island, NY) supplemented with 10% fetal bovine serum (FBS), 1% penicillin.^60^

## ASSOCIATED CONTENT

### Supporting Information

Supporting Information.pdf

## AUTHOR INFORMATION

### Author Contributions

*Conceptualiation: MH; Methodology and Data Analysis: all authors; Writing: PK, TN and MH; Project Administration: MH; Funding Acquisition: MH*

## ACKNOWLEDGMENT

MH acknowledges research grants by the Camille & Henry Dreyfus Foundation and the National Institutes of Health (NIH-NIMHD grant 8-G-12-MD007599). MPK, TN, SJ acknowledge support from the CUNY Graduate Center Graduate Fellowship program. MPK also acknowledges support form NSF-IGERT to Hunter College. Jessica Kuppan acknowledges support from the NIH-MARC program under award number T34GM007823.

## ABBREVIATIONS

VLP: Virus-Like Particle
ROMP: Ring-Opening Metathesis Polymerization
P22: Bacteriophage P22
CP: P22 coat protein
SP: P22 scaffold protein
GFP: green fluorescent protein
BBB: blood-brain barrier
DLS: dynamic light scattering
TEM: transmission electron microscopy
PDI: polydispersity index

## REFERENCES

(1) Vetter, I.; Lewis, R. J. Therapeutic Potential of Cone Snail Venom Peptides (Conopeptides). Curr. Top. Med. Chem. 2012, 12 (14), 1546–1552.

(2) Netirojjanakul, C.; Miranda, L. P. Progress and Challenges in the Optimization of Toxin Peptides for Development as Pain Therapeutics. Curr. Opin. Chem. Biol. 2017, 38, 70–79. https://doi.org/10.1016/j.cbpa.2017.03.004.

(3) Akcan, M.; Stroud, M. R.; Hansen, S. J.; Clark, R. J.; Daly, N. L.; Craik, D. J.; Olson, J. M. Chemical Re-Engineering of Chlorotoxin Improves Bioconjugation Properties for Tumor Imaging and Targeted Therapy. J. Med. Chem. 2011, 54 (3), 782–787. https://doi.org/10.1021/jm101018r.

(4) Lau, J. L.; Dunn, M. K. Therapeutic Peptides: Historical Perspectives, Current Development Trends, and Future Directions. Bioorganic Med. Chem. 2018, 26 (10), 2700–2707. https://doi.org/10.1016/j.bmc.2017.06.052.

(5) Napolitano, T.; Holford, M. Breakthroughs in Venom Peptide Screening Methods to Advance Future Drug Discovery. Protein Pept. Lett. 2018, 25 (12), 1137–1148. https://doi.org/10.2174/0929866525666181101103047.

(6) Angell, Y.; Holford, M.; Moos, W. H. Building on Success: A Bright Future for Peptide Therapeutics. Protein Pept. Lett. 2018. https://doi.org/10.2174/0929866525666181114155542.

(7) Uhlig, T.; Kyprianou, T.; Giancarlo, F.; Alberto, C.; Heiligers, D.; Hills, D.; Ribes, X.; Verhaert, P. The Emergence of Peptides in the Pharmaceutical Business?: From Exploration to Exploitation. EUPROT 2014, 4, 58–69. https://doi.org/10.1016/j.euprot.2014.05.003.

(8) Fosgerau, K.; Hoffmann, T. Peptide Therapeutics: Current Status and Future Directions. Drug Discov. Today 2015, 20 (1), 122–128. https://doi.org/10.1016/j.drudis.2014.10.003.

(9) Abbott, N. J.; Patabendige, A. A. K.; Dolman, D. E. M.; Yusof, S. R.; Begley, D. J. Structure and Function of the Blood-Brain Barrier. Neurobiol. Dis. 2010, 37 (1), 13–25. https://doi.org/10.1016/j.nbd.2009.07.030.

(10) Miljanich, G. Ziconotide: Neuronal Calcium Channel Blocker for Treating Severe Chronic Pain. Curr. Med. Chem. 2004, 11 (23), 3029–3040. https://doi.org/10.2174/0929867043363884.

(11) Atanassoff, P. G.; Hartmannsgruber, M. W. B.; Thrasher, J.; Wermeling, D.; Longton, W.; Gaeta, R.; Singh, T.; Mayo, M.; McGuire, D.; Luther, R. L. Ziconotide, a New N-Type Calcium Channel Blocker, Administered Intratehcally for Acute Postoperative Pain. Reg. Anesth. Pain Med. 2000, 25, 274–278.

(12) Kowalski, P. S.; Rudra, A.; Miao, L.; Anderson, D. G. Delivering the Messenger: Advances in Technologies for Therapeutic MRNA Delivery. Mol. Ther. 2019, 27 (4), 710–728. https://doi.org/10.1016/j.ymthe.2019.02.012.

(13) Hassett, K. J.; Benenato, K. E.; Jacquinet, E.; Lee, A.; Woods, A.; Yuzhakov, O.; Himansu, S.; Deterling, J.; Geilich, B. M.; Ketova, T.; Mihai, C.; Lynn, A.; McFadyen, I.; Moore, M. J.; Senn, J. J.; Stanton, M. G.; Almarsson, Ö.; Ciaramella, G.; Brito, L. A. Optimization of Lipid Nanoparticles for Intramuscular Administration of MRNA Vaccines. Mol. Ther. - Nucleic Acids 2019, 15 (April), s1–11. https://doi.org/10.1016/j.omtn.2019.01.013.

(14) Thanh Le, T.; Andreadakis, Z.; Kumar, A.; Gómez Román, R.; Tollefsen, S.; Saville, M.; Mayhew, S. The COVID-19 Vaccine Development Landscape. Nat. Rev. Drug Discov. 2020, 19 (5), 305–306. https://doi.org/10.1038/d41573-020-00073-5.

(15) Lian, T.; Ho, R. J. Trends and Developments in Liposome Drug Delivery Systems. J. Pharm. Sci. 2001, 90 (6), 667–680.

(16) Fleige, E.; Quadir, M. A.; Haag, R. Stimuli-Responsive Polymeric Nanocarriers for the Controlled Transport of Active Compounds: Concepts and Applications. Adv. Drug Deliv. Rev. 2012, 64 (9), 866–884. https://doi.org/10.1016/j.addr.2012.01.020.

(17) Tamanoi, F.; Zink, J. I. Multifunctional Inorganic Nanoparticles for Imaging, Targeting, and Drug Delivery. ACS Nano 2008, 2 (5), 889–896.

(18) Huo, M.; Yuan, J.; Tao, L.; Wei, Y. Redox-Responsive Polymers for Drug Delivery: From Molecular Design to Applications. Polym. Chem. 2014, 5 (5), 1519–1528. https://doi.org/10.1039/c3py01192e.

(19) Douglas, S. M.; Bachelet, I.; Church, G. M. A Logic-Gated Nanorobot for Targeted Transport of Molecular Payloads. Science (80-.). 2012, 335 (6070), 831–834. https://doi.org/10.1126/science.1214081.

(20) Rohovie, M. J.; Nagasawa, M.; Swartz, J. R. Virus-like Particles: Next-Generation Nanoparticles for Targeted Therapeutic Delivery. Bioeng. Transl. Med. 2016, 2 (1), 43–57. https://doi.org/10.1002/btm2.10049.

(21) Anand, P.; O’Neil, A.; Lin, E.; Douglas, T.; Holford, M.; O’Neil, A.; Lin, E.; Douglas, T.; Holford, M. Tailored Delivery of Analgesic Ziconotide across a Blood Brain Barrier Model Using Viral Nanocontainers. Sci. Rep. 2015, 5, 12497.

(22) Kelly, P.; Anand, P.; Uvaydov, A.; Chakravartula, S. Developing a Dissociative Nanocontainer for Peptide Drug Delivery. 2015, 12543–12555. https://doi.org/10.3390/ijerph121012543.

(23) Johnson, J. E. Virus Particle Maturation: Insights into Elegantly Programmed Nanomachines. Curr. Opin. Struct. Biol. 2010, 20 (2), 210–216. https://doi.org/10.1016/j.sbi.2010.01.004.

(24) Lee, K. W.; Tey, B. T.; Ho, K. L.; Tan, W. S. Delivery of Chimeric Hepatitis B Core Particles into Liver Cells. J. Appl. Microbiol. 2012, 112 (1), 119–131. https://doi.org/10.1111/j.1365-2672.2011.05176.x.

(25) Van Eldijk, M. B.; Wang, J. C. Y.; Minten, I. J.; Li, C.; Zlotnick, A.; Nolte, R. J. M.; Cornelissen, J. J. L. M.; Van Hest, J. C. M. Designing Two Self-Assembly Mechanisms into One Viral Capsid Protein. J. Am. Chem. Soc. 2012, 134 (45), 18506–18509. https://doi.org/10.1021/ja308132z.

(26) Fiedler, J. D.; Brown, S. D.; Lau, J. L.; Finn, M. G. RNA-Directed Packaging of Enzymes within Virus-like Particles. Angew. Chemie - Int. Ed. 2010, 49 (50), 9648–9651. https://doi.org/10.1002/anie.201005243.

(27) Glasgow, J. E.; Capehart, S. L.; Francis, M. B.; Tullman-Ercek, D. Osmolyte-Mediated Encapsulation of Proteins inside MS2 Viral Capsids. ACS Nano 2012, 6 (10), 8658–8664. https://doi.org/10.1021/nn302183h.

(28) O’Neil, A.; Reichhardt, C.; Johnson, B.; Prevelige, P. E.; Douglas, T. Genetically Programmed in Vivo Packaging of Protein Cargo and Its Controlled Release from Bacteriophage P22. Angew. Chem. Int. Ed. Engl. 2011, 50 (32), 7425–7428. https://doi.org/10.1002/anie.201102036.

(29) Patterson, D. P.; Prevelige, P. E.; Douglas, T.; Bacteriophage, P.; Patterson, D. P.; Prevelige, P. E.; Douglas, T. Nanoreactors by Programmed Enzyme Encapsulation Inside the Capsid of the Bacteriophage P22. ACS Nano 2012, 6 (6), 5000–5009. https://doi.org/10.1021/nn300545z.

(30) Wang, J.; Fang, T.; Li, M.; Zhang, W.; Zhang, Z. P.; Zhang, X. E.; Li, F. Intracellular Delivery of Peptide Drugs Using Viral Nanoparticles of Bacteriophage P22: Covalent Loading and Cleavable Release. J. Mater. Chem. B 2018, 6 (22), 3716–3726. https://doi.org/10.1039/c8tb00186c.

(31) Tsomaia, N. Peptide Therapeutics: Targeting the Undruggable Space. Eur. J. Med. Chem. 2015, 94, 459–470. https://doi.org/10.1016/j.ejmech.2015.01.014.

(32) Binder, J. B.; Raines, R. T. Olefin Metathesis for Chemical Biology. Curr. Opin. Chem. Biol. 2008, 12 (6), 767–773. https://doi.org/10.1016/j.cbpa.2008.09.022.

(33) Brendgen, T.; Fahlbusch, T.; Frank, M.; Schühle, D. T.; Seßler, M.; Schatz, J. Metathesis in Pure Water Mediated by Supramolecular Additives. Adv. Synth. Catal. 2009, 351 (3), 303–307. https://doi.org/10.1002/adsc.200800637.

(34) Hermanson, G. T. Bioconjugate Techniques.

(35) Binder, J. B.; Raines, R. T. Olefin Metathesis for Chemical Biology. Curr. Opin. Chem. Biol. 2008, 12 (6), 767–773. https://doi.org/10.1016/j.cbpa.2008.09.022.

(36) Hoveyda, A. H.; Zhugralin, A. R. The Remarkable Metal-Catalysed Olefin Metathesis Reaction. Nature 2007, 450 (7167), 243–251. https://doi.org/10.1038/nature06351.

(37) Sutthasupa, S.; Shiotsuki, M.; Sanda, F. Recent Advances in Ring-Opening Metathesis Polymerization, and Application to Synthesis of Functional Materials. Polym. J. 2010, 42 (12), 905–915. https://doi.org/10.1038/pj.2010.94.

(38) Bielawski, C. W.; Grubbs, R. H. Living Ring-Opening Metathesis Polymerization. Prog. Polym. Sci. 2007, 32 (1), 1–29. https://doi.org/10.1016/j.progpolymsci.2006.08.006.

(39) Ben-Asuly, A.; Aharoni, A.; Diesendruck, C. E.; Vidavsky, Y.; Goldberg, I.; Straub, B. F.; Lemcoff, N. G. Photoactivation of Ruthenium Olefin Metathesis Initiators. Organometallics 2009, 28 (16), 4652–4655. https://doi.org/10.1021/om9004302.

(40) Szczepaniak, G.; Kosiński, K.; Grela, K. Towards “Cleaner” Olefin Metathesis: Tailoring the NHC Ligand of Second Generation Ruthenium Catalysts to Afford Auxiliary Traits. Green Chem. 2014, 16 (10), 4474–4492. https://doi.org/10.1039/c4gc00705k.

(41) Schrodi, Y.; Pederson, R. L. Evolution and Applications of Second-Generation Ruthenium Olefin Metathesis Catalysts. 45–52.

(42) Ogba, O. M.; Warner, N. C.; O’Leary, D. J.; Grubbs, R. H. Recent Advances in Ruthenium-Based Olefin Metathesis. Chem. Soc. Rev. 2018, 47 (12), 4510–4544. https://doi.org/10.1039/c8cs00027a.

(43) Hong, S. H.; Grubbs, R. H. Highly Active Water-Soluble Olefin Metathesis Catalyst. J. Am. Chem. Soc. 2006, 128 (11), 3508–3509. https://doi.org/10.1021/ja058451c.

(44) Skowerski, K.; Szczepaniak, G.; Wierzbicka, C.; Gułajski, ł.; Bieniek, M.; Grela, K. tighly Active Catalysts for Olefin Metathesis in Water. Catal. Sci. Technol. 2012, 2 (12), 2424–2427. https://doi.org/10.1039/c2cy20320k.

(45) Skowerski, K.; Szczepaniak, G.; Wierzbicka, C.; Gułajski, ł.; Bieniek, M.; Grela, K. Highly Active Catalysts for Olefin Metathesis in Water. Catal. Sci. Technol. 2012. https://doi.org/10.1039/c2cy20320k.

(46) McCoy, K.; Selivanovitch, E.; Luque, D.; Lee, B.; Edwards, E.; Castón, J. R.; Douglas, T. Cargo Retention inside P22 Virus-Like Particles. Biomacromolecules 2018, 19 (9), 3738–3746. https://doi.org/10.1021/acs.biomac.8b00867.

(47) Goodman, L. S.; Brunton, L. L.; Chabner, B.; Knollmann, B. C. Goodman & Gilman’s Pharmacological Basis of Therapeutics, 12th ed.; McGraw-Hill: New York, NY, 2011.

(48) Parent, K. N.; Zlotnick, A.; Teschke, C. M. Quantitative Analysis of Multi-Component Spherical Virus Assembly: Scaffolding Protein Contributes to the Global Stability of Phage P22 Procapsids. J. Mol. Biol. 2006, 359 (4), 1097–1106. https://doi.org/10.1016/j.jmb.2006.03.068.

(49) Zlotnick, A.; Suhanovsky, M. M.; Teschke, C. M. The Energetic Contributions of Scaffolding and Coat Proteins to the Assembly of Bacteriophage Procapsids. Virology 2012, 428 (1), 64–69. https://doi.org/10.1016/j.virol.2012.03.017.

(50) Walker, R.; Conrad, R. M.; Grubbs, R. H. The Living ROMP of Trans-Cyclooctene. Macromolecules 2009, 2 (3), 599–605. https://doi.org/10.1021/ma801693q.

(51) Kelly, P.; Anand, P.; Uvaydov, A.; Chakravartula, S.; Sherpa, C.; Pires, E.; O’Neil, A.; Douglas, T.; Holford, M. Developing a Dissociative Nanocontainer for Peptide Drug Delivery. Int. J. Environ. Res. Public Health 2015, 12 (10), 12543–12555. https://doi.org/10.3390/ijerph121012543.

(52) Manchester, M.; Singh, P. Virus-Based Nanoparticles (VNPs): Platform Technologies for Diagnostic Imaging. Adv. Drug. Deliv. Rev. 2006, 58 (14), 1505–1522.

(53) Koudelka, K. J.; Pitek, A. S.; Manchester, M.; Steinmetz, N. F. Virus-Based Nanoparticles as Versatile Nanomachines. Annu. Rev. Virol. 2015, 2 (1), 379–401. https://doi.org/10.1146/annurev-virology-100114-055141.

(54) Kaiser, C. R.; Flenniken, M. L.; Gillitzer, E.; Harmsen, A. L.; Harmsen, A. G.; Jutila, M. A.; Douglas, T.; Young, M. J. Biodistribution Studies of Protein Cage Nanoparticles Demonstrate Broad Tissue Distribution and Rapid Clearance in Vivo. Int. J. Nanomedicine 2007, 2 (4), 715–733.

(55) Gagnon, Z. E.; Newkirk, C.; Hicks, S. Impact of Platinum Group Metals on the Environment: A Toxicological, Genotoxic and Analytical Chemistry Study. J. Environ. Sci. Heal. Part A Toxic/Hazardous Subst. Environ. Eng. 2006, 411 (3), 397–414.

(56) Patel, P. R.; Kiser, R. C.; Lu, Y. Y.; Fong, E.; Ho, W. C.; Tirrell, D. A.; Grubbs, R. H. Synthesis and Cell Adhesive Properties of Linear and Cyclic RGD Functionalized Polynorbornene Thin Films. Biomacromolecules 2012, 13 (8), 2546–2553. https://doi.org/10.1021/bm300795y.

(57) France, M. B.; Uffelman, E. S. Ring-Opening Metatheiss with a Well-Defined Ruthenium Carbene Complex An Experiment for the Undergraduate Inorganic or Polymer Laboratory. J. Chem. Educ. 1999, 76 (May), 661–665. https://doi.org/10.1021/ed076p661.

(58) Lu, H.; Wang, J.; Wang, T.; Zhong, J.; Bao, Y.; Hao, H. Recent Progress on Nanostructures for Drug Delivery Applications. J. Nanomater. 2016, 2016. https://doi.org/10.1155/2016/5762431.

(59) Taylor, M. T.; Nelson, J. E.; Suero, M. G.; Gaunt, M. J. A Protein Functionalization Platform Based on Selective Reactions at Methionine Residues. Nature 2018, 562 (7728), 563–568. https://doi.org/10.1038/s41586-018-0608-y.

(60) Anand, P.; Filipenko, P.; Huaman, J.; Lyudmer, M.; Hossain, M.; Santamaria, C.; Huang, K.; Ogunwobi, O.; Holford, M. Antitumor Effects of Tv1 Venom Peptide in Liver Cancer. bioRxiv 2019, 2, 518340. https://doi.org/10.1101/518340.

