## Supporting Information for "Induced Disassembly of a Virus-Like Particle under Physiological Conditions for Venom Peptide Delivery"

Figure S1. DNA Sequence of P22 $His_6$GFP-SP fusion protein

Figure S2. DNA Sequence of P22 Coat Protein

Figure S3. Protein Sequence of P22 $His_6$GFP-SP fusion protein

Figure S4. Protein Sequence of P22 Coat Protein

Figure S5. ESI-MS spectrum P22 Coat Protein control (nonfunctionalized)

Figure S6. Conjugation of Norbornene-COOH to P22 coat protein (MS-ESI data)

Figure S7. P22_$His_6$GFPSPCP Plasmid Map

Figure S1. DNA Sequence of P22 $His_6$GFP-SP fusion protein.

ATGCATCATCACCATCACCACAAGGGGGTGAAGGAAGTAATGAAGATCAGTCTGGAGATGGACT
GCACTGTTAACGGCGACAAATTTAAGATCATTGGGGATGGAACAGGAGAACCTTACGAAGGAAC
ACAGACTTTACATCTTACAGAGAAGGAAGGCAAGCCTCTGACGTTTTCTTTCGATGTATTGACA
CCAGCATTTCAGTATGGAAACCGTACATTCACCAAATACCCAGGCAATATACCAGACTTTTTCA
AGCAGACCGTTTCTGGTGGCGGGTATACCTGGGAGCGAAAAATGACTTATGAAGACGGGGGCAT
AAGTAACGTCCGAAGCGACATCAGTGTGAAAGGTGACTCTTTCTACTATAAGATTCACTTCACT
GGCGAGTTTCCTCCTCATGGTCCAGTGATGCAGAGGAAGACAGTAAAATGGGAGCCATCCACTG
AAGTAATGTATGTTGACGACAAGAGTGACGGTGTGCTGAAGGGAGATGTCAACATGGCTCTGTT
GCTTAAAGATGGCCGCCATTTGAGAGTTGACTTTAACACTTCTTACATACCCAAGAAGAAGGTC
GAGAATATGCCTGACTACCATTTTATAGACCACCGCATTGAGATTCTGGGCAACCCAGAAGACA
AGCCGGTCAAGCTGTACGAGTGTGCTGTAGCTCGCTATTCTCTGCTGCCTGAGAAGAACAAGGG
GCTCCCATGGCTGGTGCCGCGCGGCAGCTGTCGCAGCAATGCCGTAGCAGAACAGGGCCGCAAG
ACTCAGGAGTTTACCCAGCAATCAGCGCAATACGTCGAAGCTGCCCGCAAACACTATGACGCGG
CGGAAAAGCTCAACATCCCTGACTATCAGGAGAAAGAAGACGCATTTATGCAACTGGTTCCGCC
TGCGGTTGGGGCCGACATTATGCGCCTGTTCCCGGAAAAGTCCGCCGCGCTCATGTATCACCTG
GGGGCAAACCCGGAGAAAGCCCGCCAGTTACTGGCGATGGATGGGCAGTCCGCGCTGATTGAAC
TCACTCGACTATCCGAACGCTCTCTCAAGCCTCGCGGTAAACAAATCTCTTCCGCTCCCCATGC
TGACCAGCCTATTACCGGTGATGTCAGCGCAGCAAATAAAGATGCCATTCGTAAACAAATGGAT
GCTGCTGCGAGCAAGGGAGATGTGGAAACCTACCGCAAGCTAAAGGCAAAACTTAAAGGAATCC
GATAA

Figure S2. DNA Sequence of P22 Coat Protein.

ATGGCTTTGAACGAAGGTCAAATTGTTACACTGGCGGTAGATGAAATCATCGAAACCATCTCCG
CAATCACTCCAATGGCGCAGAAAGCCAAGAAATACACCCCGCCTGCTGCTTCTATGCAGCGCTC
CAGCAATACCATCTGGATGCCTGTAGAGCAAGAGTCACCCACTCAGGAGGGCTGGGATTTAACT
GATAAAGCGACAGGGTTACTGGAACTTAACGTCGCGGTAAACATGGGAGAGCCGGATAACGACT
TCTTCCAGTTGCGTGCTGATGACTTGCGAGACGAAACTGCGTATCGTCGCCGCATCCAGTCTGC
CGCTCGCAAGCTGGCGAACAACGTTGAGTTGTGCGTCGCAAACATGGCCGCCGAGATGGGTTCG
CTGGTTATCACCTCCCCTGATGCCATCGGCACTAATACCGCAGACGCCTGGAACTTTGTGGCCG
ACGCAGAAGAAATCATGTTCTCCCGCGAACTTAACCGCGACATGGGGACATCGTACTTCTTCAA
CCCTCAGGACTACAAAAAAGCGGGTTACGACCTGAAGAAGCGTGACATCTTCGGGCGTATTCCT
GAAGAAGCATACCGAGATGGCACCATTCAGCGTCAGGTCGCTGGCTTCGATGATGTCCTGCGCT
CTCCGAAACTTCCTGTGCTGACCAAATCCACCGCAACTGGCATCACTGTATCCGGTGCGCAGTC
CTTCAAGCCTGTCGCATGGCAACTGGATAACGATGGCAACAAAGTTAACGTTGATAACCGTTTT
GCTACCGTCACCCTGTCTGCAACTACCGGCATGAAACGCGGCGACAAAATTTCGTTTGCTGGCG
TTAAGTTCCTTGGTCAGATGGCTAAGAACGTACTGGCTCAGGATGCGACTTTCTCCGTAGTCCG
CGTTGTTGACGGTACTCATGTTGAAATCACGCCGAAGCCGGTAGCGCTGGATGATGTTTCCCTG
TCTCCGGAGCAGCGTGCCTACGCCAACGTTAACACCTCGCTGGCTGATGCAATGGCAGTGAACA
TTCTGAACGTTAAAGACGCTCGCACTAATGTGTTCTGGGCTGACGATGCTATTCGTATCGTGTC
TCAGCCGATTCCGGCTAACCATGAACTTTTTGCAGGTATGAAAACTACCTCATTCAGCATCCCT
GATGTTGGCCTGAACGGTATCTTCGCTACGCAGGGTGATATTTCCACCCTGTCCGGCCTGTGCC
GTATTGCGCTGTGGTACGGCGTAAACGCGACACGACCGGAGGCAATCGGTGTTGGCCTGCCTGG
TCAGACTGCGTAA

Figure S3. Protein Sequence of P22 $His_6$GFP-SP fusion protein.

MHHHHHHKGVKEVMKISLEMDCTVNGDKFKIIGDGTGEPYEGTQTLHLTEKEGKPLTFSFDVLT
PAFQYGNRTFTKYPGNIPDFFKQTVSGGGYTWERKMTYEDGGISNVRSDISVKGDSFYYKIHFT
GEFPPHGPVMQRKTVKWEPSTEVMYVDDKSDGVLKGDVNMALLLKDGRHLRVDFNTSYIPKKKV
ENMPDYHFIDHRIEILGNPEDKPVKLYECAVARYSLLPEKNKGLPWLVPRGSCRSNAVAEQGRK
TQEFTQQSAQYVEAARKHYDAAEKLNIPDYQEKEDAFMQLVPPAVGADIMRLFPEKSAALMYHL
GANPEKARQLLAMDGQSALIELTRLSERSLKPRGKQISSAPHADQPITGDVSAANKDAIRKQMD
AAASKGDVETYRKLKAKLKGIR*

Figure S4. Protein Sequence of P22 Coat Protein.

MALNEGQIVTLAVDEIIETISAITPMAQKAKKYTPPAASMQRSSNTIWMPVEQESPTQEGWDLT
DKATGLLELNVAVNMGEPDNDFFQLRADDLRDETAYRRRIQSAARKLANNVELCVANMAAEMGS
LVITSPDAIGTNTADAWNFVADAEEIMFSRELNRDMGTSYFFNPQDYKKAGYDLKKRDIFGRIP
EEAYRDGTIQRQVAGFDDVLRSPKLPVLTKSTATGITVSGAQSFKPVAWQLDNDGNKVNVDNRF
ATVTLSATTGMKRGDKISFAGVKFLGQMAKNVLAQDATFSVVRVVDGTHVEITPKPVALDDVSL
SPEQRAYANVNTSLADAMAVNILNVKDARTNVFWADDAIRIVSQPIPANHELFAGMKTTSFSIP
DVGLNGIFATQGDISTLSGLCRIALWYGVNATRPEAIGVGLPGQTA*

Figure S5. ESI-MS spectrum P22 Coat Protein control (nonfunctionalized).
M.W. = 46623.9 Da (calculated 46622.7 Da)

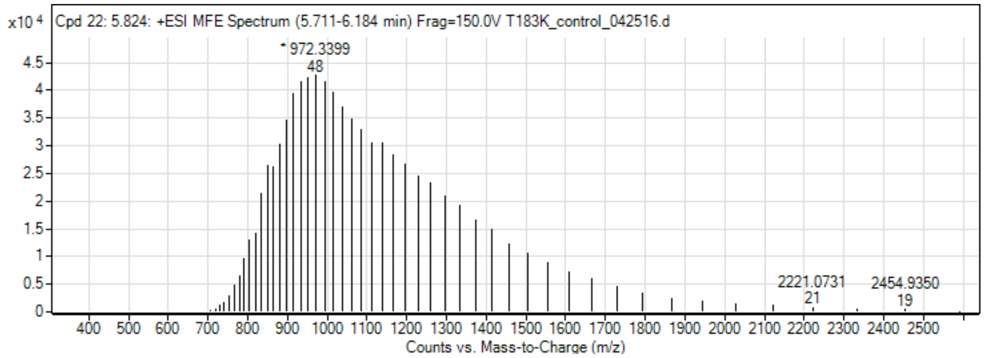

Figure S6. Conjugation of Norbornene-COOH to P22 coat protein (MS-ESI data)

| Compound ID | Mass (Da) | Vol | Vol% Of Observed Coat Proteins | No. of conjugated Norbornenes (Norbornene M.W. = 120) |
|---|---|---|---|---|
| 1 | 47226.00 | 214651 | 36.8 | 5 |
| 2 | 47106.61 | 180238 | 30.9 | 4 |
| 3 | 47344.52 | 91391 | 15.6 | 6 |
| 4 | 47465.05 | 42537 | 7.3 | 7 |
| 5 | 47584.17 | 32840 | 5.6 | 8 |
| 6 | 46985.02 | 22404 | 3.8 | 3 |

Figure S7. P22_His$_6$GFPSPCP Plasmid Map

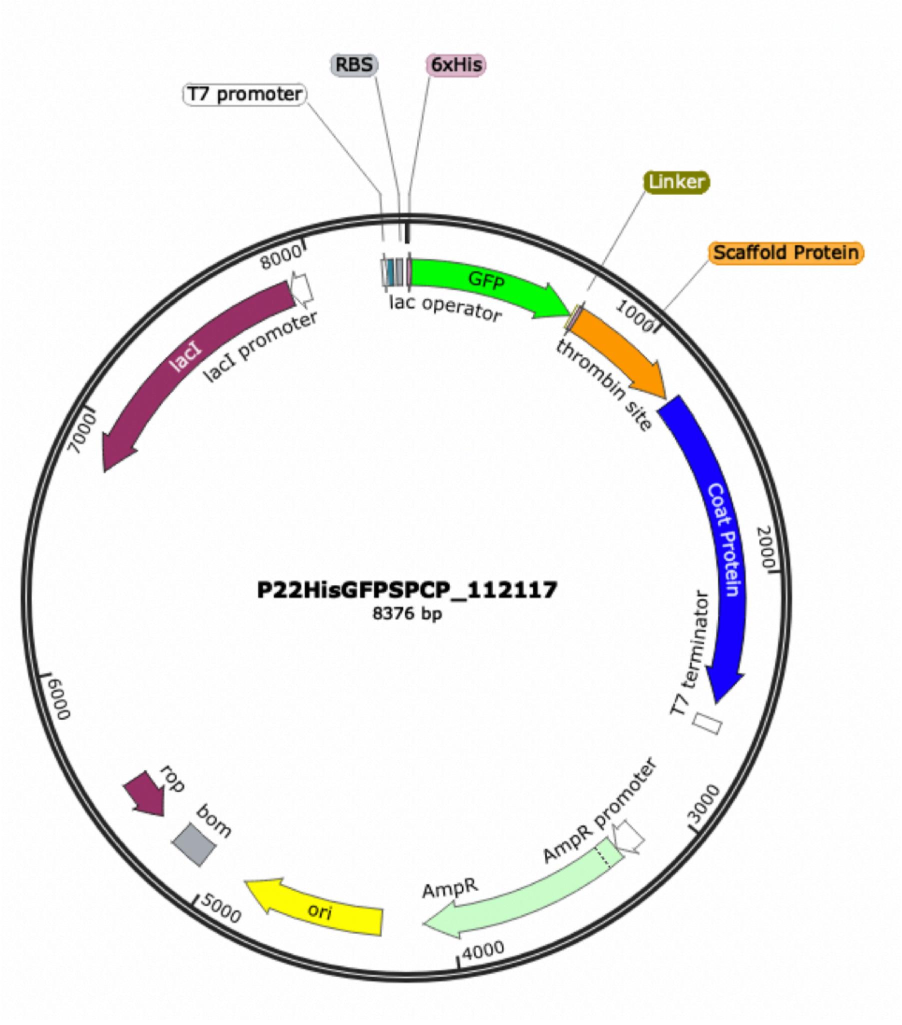
